# Delay Differential Equation (DDE) Modeling of CAR-T Cellular Kinetics: Application to BCMA-Targeted (Ide-cel, Orva-cel) and CD19-Targeted (Liso-cel) Therapies

**DOI:** 10.64898/2026.03.01.708830

**Authors:** Yiming Cheng, Yan Li

**Author notes:** Corresponding Author: Yan Li, PhD, Clinical Pharmacology & Pharmacometrics, Bristol Myers Squibb, 556 Morris Avenue, Summit, NJ 07901. Disclosures: Y.C. and Y.L. are employees and hold equity ownership in Bristol Myers Squibb. **Data Sharing**: The datasets are available from the corresponding author upon request.

## Abstract

Chimeric antigen receptor (CAR) T-cell therapies undergo rapid in vivo expansion followed by contraction and variable long-term persistence after a single infusion, yielding cellular kinetic (CK) profiles that differ fundamentally from conventional small-molecule and biologic pharmacokinetics. Piecewise, phase-based CK models are widely used, but discontinuous switching and constant expansion assumptions can create numerical instability around the transition window and bias the characterization of early expansion and near-peak behavior. Building on our prior saturable expansion framework (Vmax/Km), we advanced CAR-T CK modeling by introducing (i) smooth S-shaped gating to replace discontinuous phase switching and enable continuous time-varying expansion dynamics, and (ii) delay differential equation (DDE) components to evaluate whether longitudinal data support explicit lags in downstream biological processes. Data were pooled from three CAR-T trials (TRANSCEND, KarMMa-3, and EVOLVE) spanning two BCMA-targeted products (ide-cel, orva-cel) and one CD19-targeted product (liso-cel). Models were estimated in Monolix using SAEM with importance sampling for final likelihood evaluation. Model selection relied on likelihood-based criteria (AIC, BIC, and Monolix BICc) and diagnostic assessments. Relative to a constant-expansion baseline, saturable expansion improved fit and reduced systematic model misspecification at high transgene levels (e.g., qPCR transgene copies/µg). Among multiple DDE placements evaluated, the data most strongly supported a delay on effector-to-memory conversion; delays in effector-like and/or memory-like decay were not favored. Simulations indicated that the conversion delay primarily modulated the timing and magnitude of the memory-like trajectory, with minimal impact on total-cell trajectories during the expansion phase at the evaluated scale. In a shared covariate framework with product as a categorical effect, BCMA-targeted products exhibited higher baseline levels and expansion capacity than liso-cel, with stronger evidence for slower effector-like decay for orva-cel than ide-cel. Overall, smooth-gated saturable expansion with DDE-based delayed conversion provides a parsimonious, biologically motivated framework for CAR-T CK and supports cross-product comparisons under harmonized structural assumptions.

## Introduction

Chimeric antigen receptor (CAR) T-cell therapies are a distinct modality in which the “drug” is a living cell population that can undergo rapid in vivo expansion, contraction, and long-term persistence after a single infusion. ^1–5^ Consequently, CAR-T cellular kinetics (CK) differ fundamentally from those of small molecules and monoclonal antibodies. ^3^ These features motivate semi-mechanistic pharmacometric models that (i) represent key biological programs with a parsimonious and interpretable structure, (ii) remain identifiable under typical clinical sampling, and (iii) enable consistent inference across studies and products. ^6–11^

A widely used baseline framework for CAR-T CK is a piecewise, phase-based model motivated by classical immune-response theory. ^11, 12^ In this construct, an activated/effector-like compartment expands for a finite period and then transitions into contraction/persistence. After the transition, activated cells may be lost rapidly and/or convert into a longer-lived memory-like compartment, and both compartments subsequently decline via first-order decay. ^3, 6, 8, 9, 11, 13^ Practical experience, however, highlights limitations of strictly piecewise, shared-transition models. Abrupt switching imposes instantaneous rate changes that can generate numerical artifacts during estimation and simulation, particularly when data are noisy or sparse around the transition window. ^14^ Moreover, forcing distinct biological processes to share a single synchronized transition time is physiologically restrictive: attenuation of expansion, initiation of differentiation/conversion, and emergence of persistence programs need not occur simultaneously or over the same time window. Finally, assuming constant (non-saturable) expansion can be too rigid to reproduce rapid early growth followed by gradual slowing as cell numbers increase, leading to systematic lack-of-fit at early time points and at the upper end of observed concentrations.

In our prior work, we addressed the constant-expansion limitation by replacing the expansion term with a saturable formulation (Vmax/Km), which increased flexibility during early expansion and around peak and improved empirical fit while preserving an interpretable semi-mechanistic structure suitable for population analysis. ^8, 10, 13^ Building on that foundation, the current work advances the framework in two directions. First, we replace discontinuous piecewise switching with smooth S-shaped gating functions to enable continuous, time-varying transitions and relax the assumption of an instantaneous phase boundary. Second, we evaluate whether CAR-T CK data support an additional mechanistic feature: explicit time delays in downstream biological processes.

Biologically, delays are expected because many cellular processes unfold over time. ^15^ In T-cell biology, temporal thresholds between stimulation and downstream outcomes have been described, and fate decisions related to effector versus memory programs may begin during expansion but only manifest in measurable compartment dynamics after a lag. ^16^ In CAR-T systems, such delays are plausibly relevant to the emergence and accumulation of memory-like phenotypes that may be numerically small relative to the expansion-phase effector-like pool yet central to longer-term persistence. ^17, 18^

From a modeling standpoint, delay differential equations (DDEs) provide a principled way to represent such lags by allowing the rate of change of a state to depend explicitly on its past values. ^19, 20^ This is mechanistically distinct from an observation-level lag time, which shifts the time at which an observation reflects the underlying state (e.g., *y(t) = f(x(t-T_lag_))*) without altering the governing dynamics. In contrast, DDEs modify the system dynamics directly (e.g., *dx/dt = g(x(t), x(t-τ))*), thereby changing trajectories of both observed and latent compartments. DDEs have been used broadly in immunodynamics and tumor–immune modeling to represent delayed proliferation, differentiation, transport, or intracellular processes, and modern NLME software (including Monolix) enables practical implementation and estimation of DDE-based models in a population context. ^19, 21–23^

Accordingly, we extended the saturable expansion framework by incorporating smooth gating and DDE components and systematically evaluating alternative delay placements. Specifically, we tested whether longitudinal CAR-T CK data support a delay in conversion from effector-like to memory-like cells and, as competing hypotheses, whether delays in effector-like decay and/or memory-like decay are warranted. The objective of this work is to determine whether CK data from three CAR-T therapies—two BCMA-targeted products (ide-cel and orva-cel) and one CD19-targeted product (liso-cel)—support delayed dependence, and, if so, to identify which underlying biological process best explains the observed longitudinal kinetics.

## Methods

### Clinical Study Data

Data were pooled from three clinical studies of CAR-T therapy—TRANSCEND, KarMMa-3, and EVOLVE—as described previously. ^9, 10, 13^ In brief, baseline demographic and disease characteristics were broadly comparable across studies, supporting analysis under a generally similar patient population framework.

All studies were conducted in accordance with the Declaration of Helsinki, ICH Good Clinical Practice (GCP) guidelines, and applicable local regulatory requirements. Protocols and amendments were approved by institutional review boards or independent ethics committees at each participating site, and all participants provided written informed consent prior to any study-specific procedures.

### Software and Estimation Methods

Data assembly and post-processing were performed in R/RStudio (R v4.1.3; Posit Software, Boston, MA, USA). Model development and parameter estimation were conducted in Monolix® (2024R1; Lixoft SAS, Antony, France). Population parameters were estimated using the stochastic approximation expectation–maximization (SAEM) algorithm, and the final log-likelihood and objective function value (OFV) were computed using importance sampling.

Model selection was guided by the log-likelihood, Akaike information criterion (AIC), Bayesian information criterion (BIC), and Monolix BICc (corrected BIC), which adds a small-sample penalty based on the number of subjects and the effective number of estimated parameters.

All candidate models were implemented in the Mlxtran language. Both ordinary differential equation (ODE) and delay differential equation (DDE) systems were evaluated; DDE models were specified using the built-in delay() function and solved using Monolix’s dedicated DDE solver. Inter-individual variability was modeled using exponential random effects (η ∼ N(0, Ω)), implying lognormal distributions for individual parameters. Residual error was modeled as additive on the log scale (normally distributed with mean 0 and variance σ²). BLQ observations were handled using the M3 likelihood-based censoring method. In covariate testing, categorical covariates were implemented on the log-parameter scale using indicator variables (e.g., *log(θ_i_) = log(θ_pop_) β I_group_ + η_i_*); therefore, represents a log-multiplicative effect and the corresponding fold-change on the original parameter scale is exp β.

### Population cellular kinetic structure model

Three semi-mechanistic population cellular kinetic (CK) model structures were evaluated in parallel. The reference structure was a widely used piecewise, shared-transition model motivated by the theoretical work of De Boer and Perelson on vigorous immune responses. ^24^ In this framework, CAR-T kinetics are represented by an initial expansion phase followed by a biphasic contraction/persistence phase. During early expansion, activated/effector-like CAR-T cells proliferate rapidly for a finite period under a non–antigen-limited growth assumption. After a transition time (*t_max1_*), the system enters contraction, during which activated cells either undergo rapid loss or convert into a longer-lived memory-like population. Activated and memory-like compartments then decline with their respective first-order rate constants.

Let A(t) denote the activated/effector-like CAR-T cells and M(t) the memory-like CAR-T cells. The baseline piecewise model is defined as:

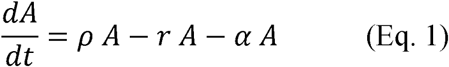

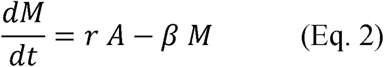

with piecewise activation of process rates governed by:

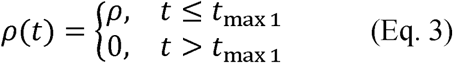

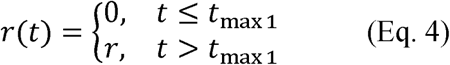

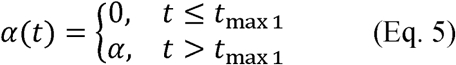

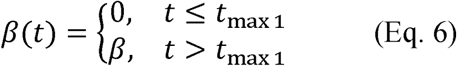

where ρ is the expansion rate constant, is the conversion rate constant from to *A* to *M*, α is the activated/effector-like decay rate constant, and is the memory-like decay rate constant.

In our prior work (Supplementary Figure 1), ^10, 13, 25^ we extended the baseline piecewise constant expansion framework by introducing saturable expansion during the expansion phase, which improved model fit, particularly at early time points. Specifically, the expansion rate for t ≤ tmax1 in Eq. 3 was replaced by:

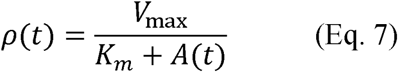

Here, % denotes the maximal activated-cell expansion rate, & is the activated-cell level at half-maximal expansion.

In the current work, we further extended the framework in two ways. First, we replaced the discontinuous piecewise switching with a continuous S-shaped gating function, allowing the expansion rate to taper gradually over time rather than switching off abruptly. Second, we incorporated delay differential equation (DDE) components to represent biologically plausible lags in downstream processes. Under this formulation, the time-varying saturable expansion rate is given by:

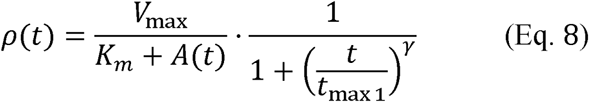

and DDE variants were specified by introducing delayed dependence in conversion and/or decay terms:

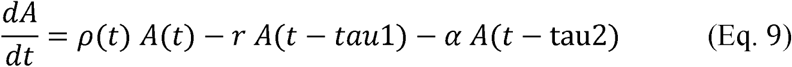

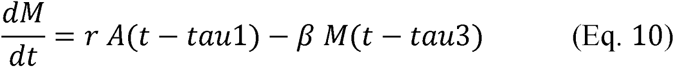

Here *t_max1_* is the characteristic time governing the decline of expansion, and γ controls the steepness of the gating transition. Under the smooth-gating saturable expansion model, *r*, α, and β were treated as time-invariant (i.e., not piecewise activated). The delay parameters tau1, tau2, and tau3 represent lags associated with effector-to-memory conversion, activated/effector-like decay, and memory-like decay, respectively. Although tau1, tau2, and tau3 were included in the general DDE formulation, inclusion of specific delay terms in the final model was determined empirically based on data support and comparative model performance.

### Simulations

Population-level simulations were conducted using the final population parameter estimates from each candidate model. Simulations were implemented in R using Simulx via the mlxR package. For each model, the time courses of activated/effector-like cells (*A*), memory-like cells (*M*), and total cells (*A + M*) were generated simultaneously. All simulation summaries and figures were produced in R.

## Results

### Model Development: Saturable Expansion and a Conversion-Delay DDE Best Describe CAR-T Cellular Kinetics

Piecewise cellular kinetic models are widely used to describe CAR-T expansion, contraction, and persistence, and the constant expansion rate (rho) structure (Model #1; constant expansion rate rho) has been applied extensively as a practical baseline model. In our previous work, we advanced this conventional framework by replacing the constant expansion term with a saturable expansion function (Vmax; Model #2, using Vmax/Km), which provided greater flexibility and improved empirical fit, particularly for the early expansion phase. In this work, we further extended the Vmax framework in two ways: (i) replacing discontinuous piecewise switching with smooth S-shaped gating functions for a time-varying expansion rate, and (ii) incorporating a delay differential equation (DDE) component to represent biologically plausible lags in downstream processes, yielding Vmax::r (Model #3; saturable expansion with a DDE applied to the conversion process r). In addition, we explored alternative DDE configurations by applying delays to different processes, including r, alpha, beta, and combinations thereof (e.g., r/alpha/beta denotes delays applied simultaneously to conversion and both decay processes).

Table 1 summarizes parameter estimates for the three base models (rho, Vmax, and Vmax::r). Relative to rho, the saturable expansion model (Vmax) estimated a later peak timing (tmax1 ≈ 8 days vs 6.5 days), consistent with improved capture of early expansion dynamics. The conversion rate constant (r) increased from rho to Vmax and remained higher in Vmax::r model. The effector-like decay parameter (alpha) was similar across models (∼0.10), whereas the DDE model (Vmax::r) estimated a modestly higher memory-like decay rate (beta: 0.009 vs 0.0067– 0.0074 in rho/Vmax). In the selected DDE model (Vmax::r), the estimated conversion delay was tau1 = 2.63 days (Table 1). Overall, fixed-effect parameters were estimated with good precision, as indicated by low relative standard errors (Table 1), and key parameters were supported by reasonably tight 2.5th–97.5th percentile intervals.

**Table 1.**
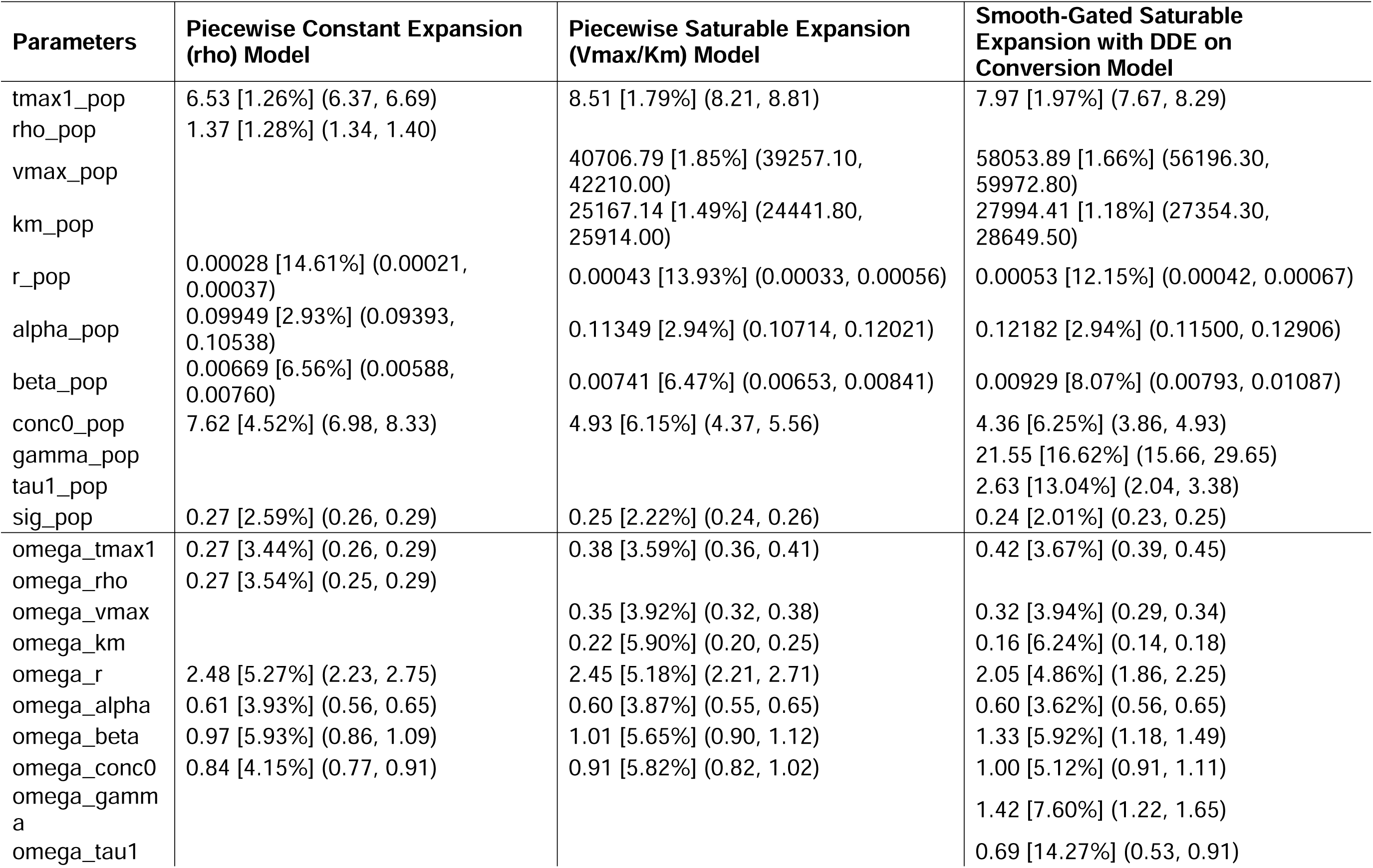
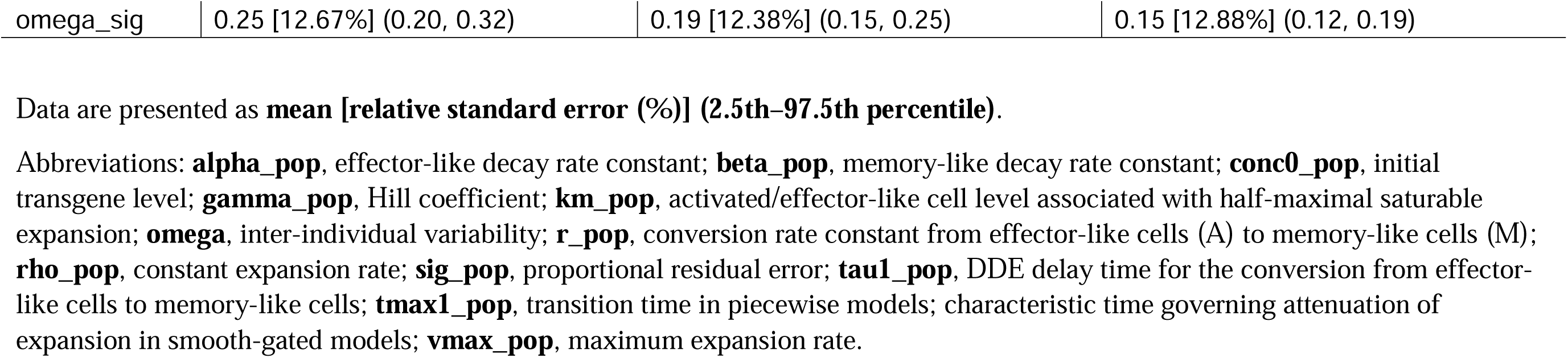
Population cellular kinetics parameter estimates for the piecewise constant, piecewise saturable, and saturable expansion with DDE-based conversion models. Data are presented as mean [relative standard error (%)] (2.5th–97.5th percentile).

Figure 1 presents ΔAIC/ΔBIC/ΔBICc/ΔOFV using the Vmax model as the reference. Consistent across criteria, the saturable expansion model substantially improved fit compared with the constant expansion model (rho), with large decreases in information criteria and OFV. Adding DDE structure further improved model performance. Among all DDE configurations tested, applying the delay to conversion only (Vmax::r) provided the largest incremental improvement, whereas adding delays to alpha and/or beta—either alone or in combination (e.g., r/alpha, r/beta, r/alpha/beta)—did not yield comparable gains. These results indicate that the data support a delay in the conversion process, while additional delays in effector-like and/or memory-like decay processes were not favored by the model-selection criteria compared with the model applying the delay to conversion only. Under Vmax::r, the conversion delay (tau1) was estimated with acceptable precision (Table 1), supporting identifiability of the delay parameter in this configuration.

**Figure 1.**
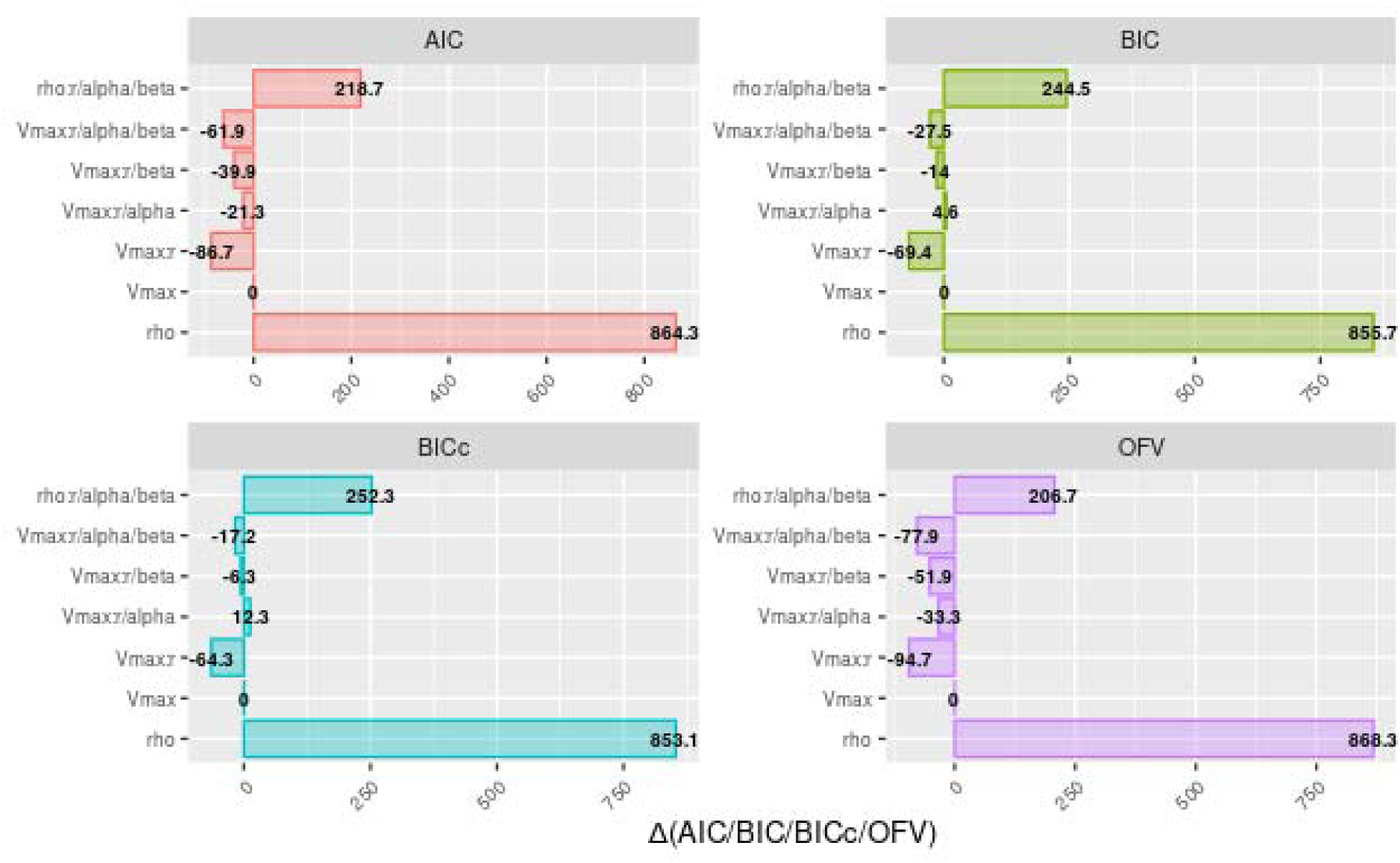
Model-selection comparison across CAR-T cellular kinetic structures: ΔAIC, ΔBIC, ΔBICc, and ΔOFV for the piecewise constant expansion (rho) and DDE variants relative to the piecewise saturable expansion reference model. Model abbreviations: rho, piecewise constant expansion; Vmax, piecewise saturable expansion (Vmax/Km); Vmax::r, smooth-gated saturable expansion with a DDE delay on conversion; Vmax::r/alpha, Vmax::r with an additional delay on effector-like decay (alpha); Vmax::r/beta, Vmax::r with an additional delay on memory-like decay (beta); Vmax::r/alpha/beta, Vmax::r with delays on conversion, alpha, and beta; rho::r/alpha/beta, piecewise constant expansion with delays on conversion, alpha, and beta. AIC, Akaike information criterion; BIC, Bayesian information criterion; BICc, corrected BIC as implemented in Monolix; OFV, objective function value.

Figure 2 shows observed vs model-predicted plots on the log scale for each candidate structure, where the red line denotes the identity line and the blue line denotes the LOESS smoother. The rho model showed systematic model misspecification, most evident at higher concentrations. The Vmax model reduced this bias, consistent with improved characterization of early expansion and the upper range of observations. Importantly, incorporation of the DDE component (Vmax::r) further improved alignment between the LOESS smoother and the identity line—particularly in the mid-range—supporting the added delay mechanism on conversion (Figure 2b). Overall, the combined evidence from information criteria (Figure 1), parameter estimates (Table 1), and diagnostics (Figure 2) supports selection of Vmax::r as the best-performing structural model among the candidates evaluated.

**Figure 2.**
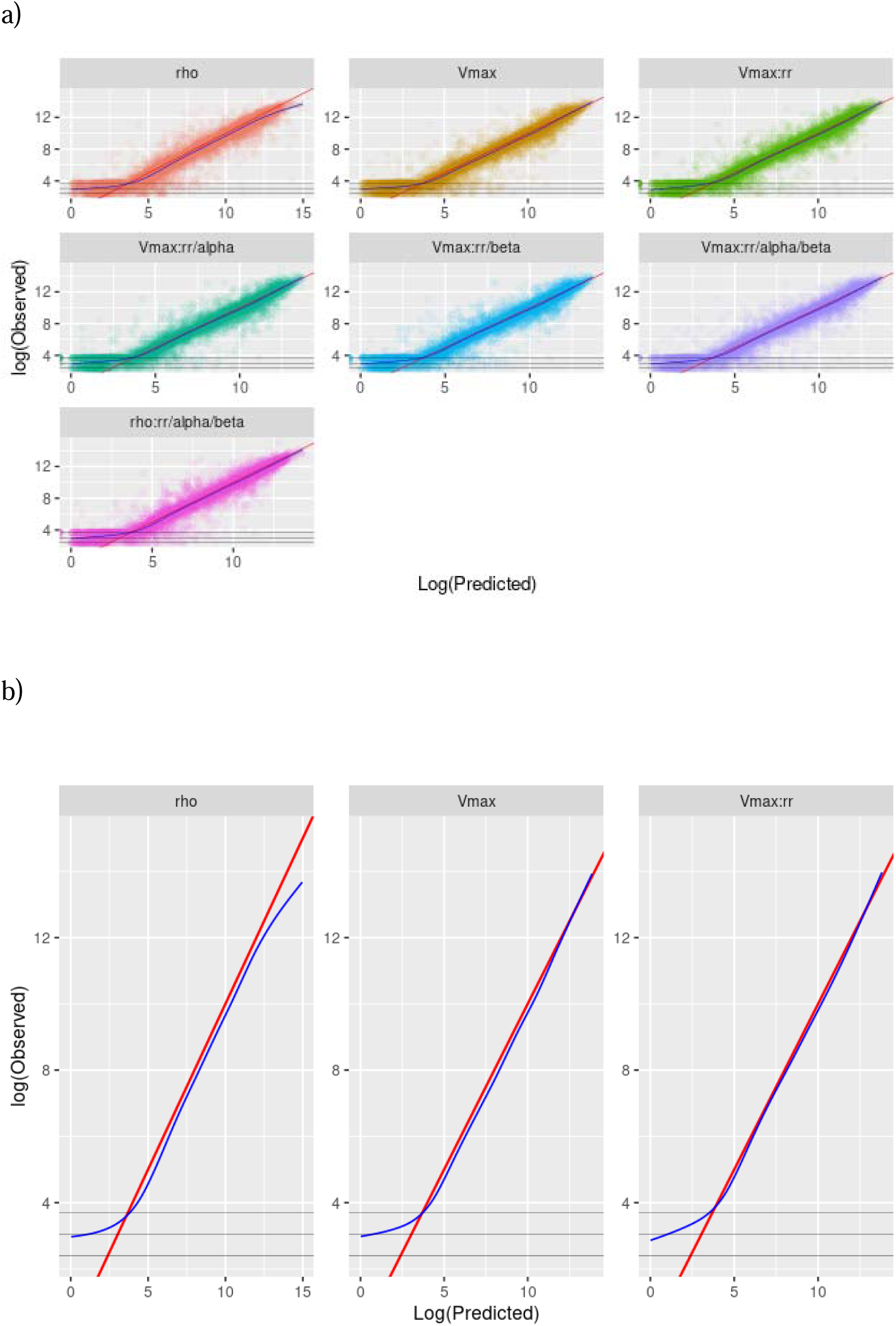
Goodness-of-fit diagnostics for candidate CAR-T cellular kinetic models: observed versus population model–predicted transgene levels. Model abbreviations: rho, piecewise constant expansion; Vmax, piecewise saturable expansion (Vmax/Km); Vmax::r, smooth-gated saturable expansion with a DDE delay on conversion; Vmax::r/alpha, Vmax::r with an additional delay on effector-like decay (alpha); Vmax::r/beta, Vmax::r with an additional delay on memory-like decay (beta); Vmax::r/alpha/beta, Vmax::r with delays on conversion, alpha, and beta; rho::r/alpha/beta, piecewise constant expansion with delays on conversion, alpha, and beta. The red line denotes the line of identity and the blue line denotes the LOESS smoother. Horizontal dotted lines indicate the assay lower limit of quantification (LLOQ) for each therapy (11, 21, and 40 copies/µg for liso-cel, orva-cel, and ide-cel, respectively). Panel (A) shows all candidate models; Panel (B) provides a focused comparison of rho, Vmax, and Vmax::r models to highlight differences in **mid-range** agreement with the line of identity.

### Simulated Effector-, Memory-, and Total-Cell Profiles Across CAR-T Cellular Kinetic Models

We evaluated multiple CAR-T cellular kinetic models, including the historical **rho** model (piecewise constant expansion), the improved **Vmax** model (piecewise saturable expansion), and the **Vmax::r** model (saturable expansion with a DDE applied to the conversion process). To compare the implications of these structures on predicted cellular kinetics, we simulated the time courses for **effector-like cells**, **memory-like cells**, and **total cells** using the final parameter estimates from each model. Figures 3a and 3b display the simulated profiles on the **log scale** (Figure 3a) and **linear scale** (Figure 3b), respectively.

**Figure 3.**
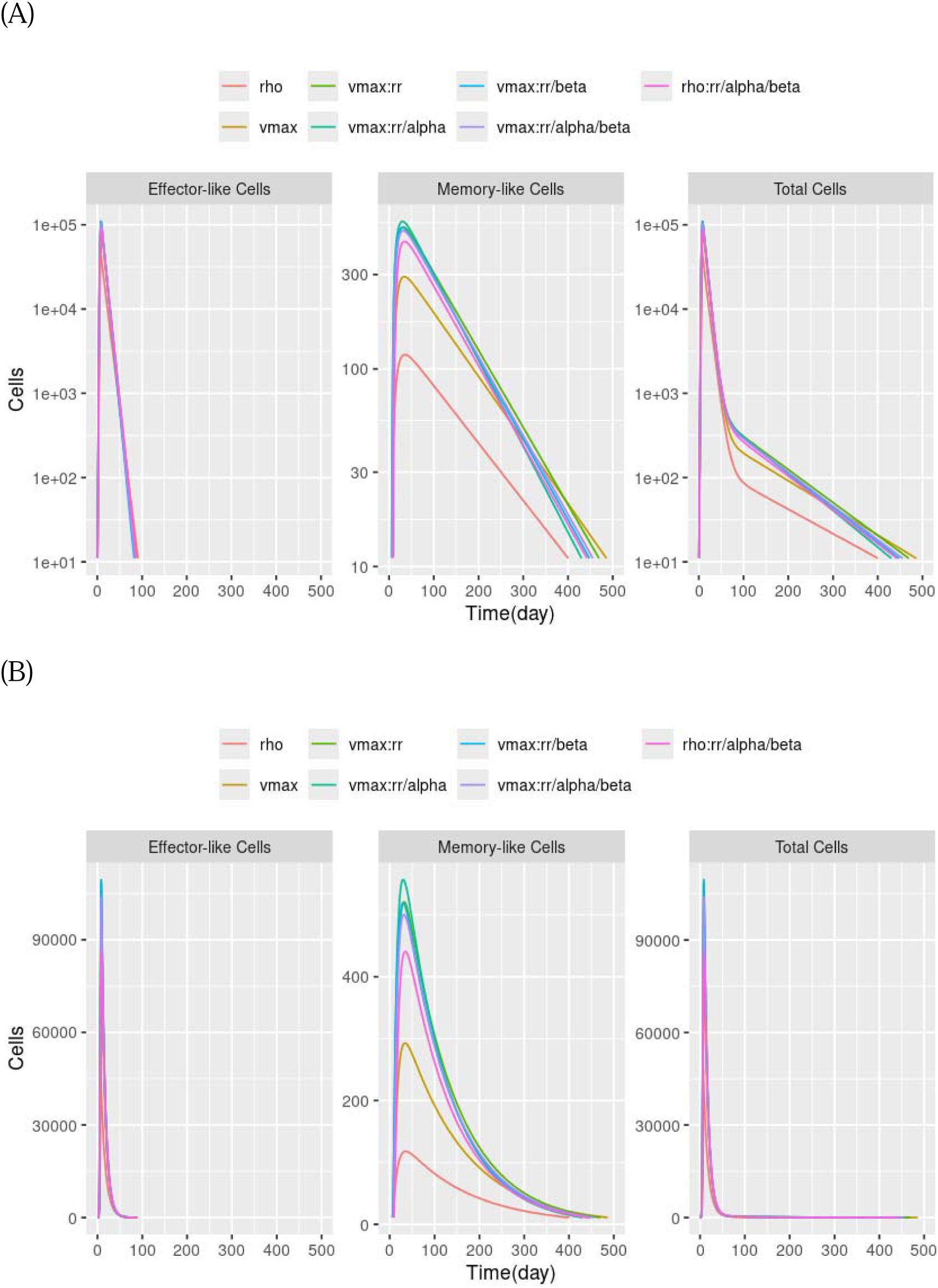
Simulated effector-like, memory-like, and total CAR-T cellular kinetics under alternative structural models. Model abbreviations: rho, piecewise constant expansion; Vmax, piecewise saturable expansion (Vmax/Km); Vmax::r, smooth-gated saturable expansion with a DDE delay on conversion; Vmax::r/alpha, Vmax::r with an additional delay on effector-like decay (alpha); Vmax::r/beta, Vmax::r with an additional delay on memory-like decay (beta); Vmax::r/alpha/beta, Vmax::r with delays on conversion, alpha, and beta; rho::r/alpha/beta, piecewise constant expansion with delays on conversion, alpha, and beta. Panels show simulations on (A) log scale and (B) linear scale.

Across all three compartments, the **rho** model produced lower predicted cell levels than the Vmax-based models, with the most pronounced separation observed during the expansion phase and around the peak on the linear-scale plots (Figure 3b). Consistent with this, the rho model showed the smallest peak magnitude in memory-like cells and consequently lower total cells.

The Vmax model increased the magnitude of the expansion-phase predictions relative to rho and yielded higher simulated profiles for both effector-like cells and total cells.

Among Vmax-based models, incorporating a DDE on conversion (Vmax::r) primarily affected the simulated memory-like cell trajectory, which was higher than Vmax over a broad time range in both the linear and log displays. The impact of Vmax::r on total cells was correspondingly visible on the log-scale plot (Figure 3a), where differences among models are more apparent at lower concentrations and later times. Overall, Figures 3a–3b show that moving from rho to Vmax and then to Vmax::r leads to progressively higher simulated cell profiles, with the largest incremental differences under Vmax::r most evident in the memory-like and total-cell trajectories.

### Sensitivity Analysis: Effect of Conversion Delay (tau1) on Simulated Effector-, Memory-, and Total-Cell Profiles

To evaluate the influence of the conversion delay parameter (tau1) in the final DDE model, we simulated effector-like, memory-like, and total-cell profiles across a range of tau1 values while holding all other parameters fixed. Figure 4 summarizes the resulting simulated trajectories.

**Figure 4.**
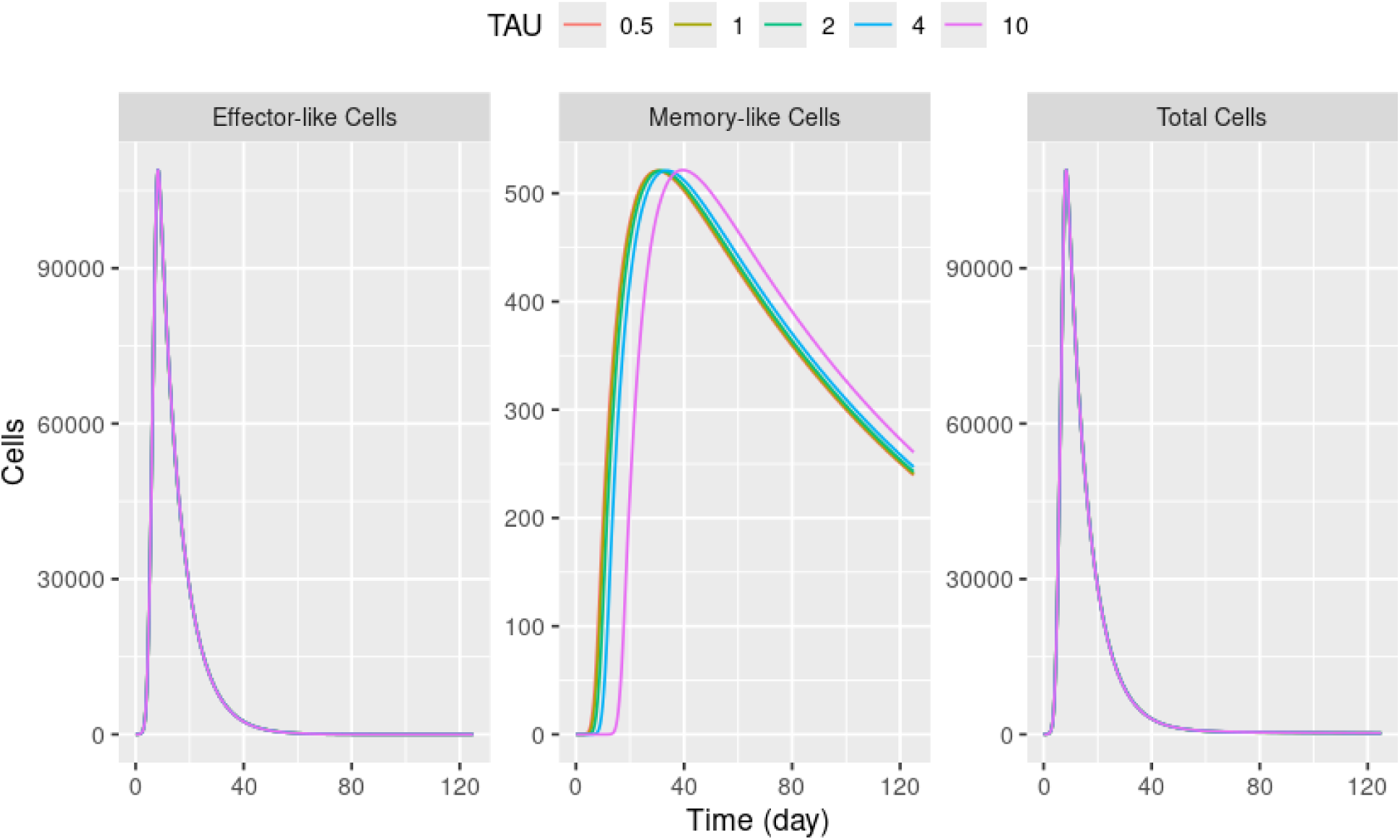
Sensitivity of simulated CAR-T cellular kinetics to the conversion delay (tau1) in the smooth-gated saturable expansion model with delayed conversion. tau1 was varied across a predefined range while all other parameters were fixed at the final population estimates; simulated trajectories are shown for effector-like cells, memory-like cells, and total cells.

Varying tau1 produced a clear and systematic effect on the memory-like cell profile, with larger tau1 values shifting the onset, rise, and peak of memory-like cells to later times. In contrast, the effector-like cell profile showed minimal sensitivity to tau1 across the tested range. The total-cell profile also exhibited little visible change with tau1 at the scale shown. This pattern is consistent with the relative magnitudes of the compartments in Figure 4: effector-like and total cells peak on the order of ∼10^5 cells, whereas memory-like cells peak on the order of ∼10^2–10^3 cells, such that changes in memory-like cells have limited impact on total cells in these simulations.

Collectively, these results indicate that tau1 is primarily informed by—and primarily influences—the predicted timing of the memory-like component, with limited impact on the effector-like and total-cell trajectories under the conditions evaluated.

### Comparison of Cellular Kinetic Parameters Across CAR-T Therapies

Because Vmax::r provided the best overall fit, and the analysis included three CAR-T products spanning two targets (BCMA: orva-cel, ide-cel; CD19: liso-cel), we compared cellular kinetic parameters across therapies under a consistent model structure. Drug product was incorporated as a categorical covariate in the final base model (Vmax::r) to assess product-specific differences while holding the structural assumptions fixed. Table 2 summarizes the covariate model estimates.

**Table 2.** Population cellular kinetics parameter estimates for the saturable expansion model with DDE-based conversion, including drug product as a categorical covariate (reference: liso-cel).

| Parameters | Mean | Relative Standard Error% | 2.5th percentile | 97.5th percentile |
| --- | --- | --- | --- | --- |
| tmax1_pop | 8.31 | 1.81 | 8.02 | 8.61 |
| r_pop | 0.0005 | 11.30 | 0.0004 | 0.0006 |
| alpha_pop | 0.1459 | 5.34 | 0.1315 | 0.1620 |
| beta_alpha_orva-cel | -0.44 | 17.23 | -0.59 | -0.29 |
| beta_alpha_ide-cel | -0.09 | 73.76 | -0.23 | 0.04 |
| beta_pop | 0.0088 | 7.26 | 0.0077 | 0.0102 |
| conc0_pop | 1.69 | 10.10 | 1.39 | 2.06 |
| beta_conc0_orva-cel | 0.43 | 38.33 | 0.11 | 0.76 |
| beta_conc0_ide-cel | 1.70 | 7.59 | 1.45 | 1.96 |
| sig_pop | 0.23 | 2.17 | 0.22 | 0.24 |
| vmax_pop | 44049.40 | 4.36 | 40448.10 | 47971.30 |
| beta_vmax_orva-cel | 0.39 | 11.03 | 0.31 | 0.47 |
| beta_vmax_ide-cel | 0.20 | 19.38 | 0.12 | 0.27 |
| km_pop | 25708.14 | 3.66 | 23928.50 | 27620.10 |
| tau1_pop | 2.50 | 18.41 | 1.76 | 3.56 |
| gamma_pop | 19.27 | 12.13 | 15.23 | 24.38 |
| omega_tmax1 | 0.38 | 3.72 | 0.35 | 0.41 |
| omega_r | 2.00 | 4.43 | 1.83 | 2.18 |
| omega_alpha | 0.59 | 4.10 | 0.55 | 0.64 |
| omega_beta | 1.28 | 5.30 | 1.15 | 1.42 |
| omega_conc0 | 0.91 | 5.19 | 0.82 | 1.01 |
| omega_sig | 0.21 | 10.66 | 0.17 | 0.25 |
| omega_vmax | 0.76 | 3.57 | 0.71 | 0.82 |
| omega_km | 0.77 | 3.80 | 0.71 | 0.83 |
| omega_tau1 | 0.81 | 38.43 | 0.41 | 1.62 |
| omega_gamma | 1.30 | 5.95 | 1.16 | 1.46 |
| corr_km_conc0 | -0.37 | 18.78 | -0.49 | -0.22 |
| corr_vmax_conc0 | -0.54 | 10.32 | -0.64 | -0.42 |
| corr_vmax_km | 0.90 | 1.27 | 0.87 | 0.92 |
| corr_tmax1_beta | 0.46 | 12.41 | 0.34 | 0.56 |

Abbreviations: **alpha\_pop**, effector-like decay rate constant; **beta\_alpha\_ide-cel**, effect of ide-cel on the alpha decay rate; **beta\_alpha\_orva-cel**, effect of orva-cel on the alpha decay rate; **beta\_conc0\_ide-cel**, effect of ide-cel on the initial transgene level; **beta\_conc0\_orva-cel**, effect of orva-cel on the initial transgene level; **beta\_pop**, memory-like decay rate constant; **beta\_vmax\_ide-cel**, effect of ide-cel on the maximum expansion rate; **beta\_vmax\_orva-cel**, effect of orva-cel on the maximum expansion rate; **conc0\_pop**, initial transgene level; **corr**, correlation coefficient between parameters (as specified by the corr\_\* entries); **gamma\_pop**, Hill coefficient; **km\_pop**, activated/effector-like cell level associated with half-maximal saturable expansion; **omega**, inter-individual variability; **r\_pop**, conversion rate constant from effector-like cells (A) to memory-like cells (M); **sig\_pop**, proportional residual error; **tau1\_pop**, DDE delay time for the conversion from effector-like cells to memory-like cells; **tmax1\_pop**, transition time in piecewise models; characteristic time governing attenuation of expansion in smooth-gated models; **vmax\_pop**, maximum expansion rate.

Most fixed-effect parameters were comparable to those from the base Vmax::r model. Notably, the covariate model yielded lower estimates of Vmax and Km than the base model, while maintaining similar estimates for key timing and rate parameters (e.g., tmax1, r, alpha, beta, and tau1). The covariate effects suggested product-level differences in baseline level, expansion capacity, and effector-like decay.

Relative to liso-cel, both BCMA-targeted products exhibited higher baseline concentrations (conc0). The estimated covariate effects were 0.43 for orva-cel and 1.70 for ide-cel, corresponding to approximately 1.54-fold and 5.47-fold increases on the original scale (exp(0.43) and exp(1.70), respectively).

In addition, the model suggested slower effector-like decay (alpha) for orva-cel relative to liso-cel (beta_alpha_orva-cel = −0.44; exp(−0.44) ≈ 0.64). For ide-cel, the estimated alpha effect was small and imprecise (beta_alpha_ide-cel = −0.09; 95% interval includes 0), indicating limited evidence for a difference relative to liso-cel. Finally, both BCMA products showed higher expansion capacity relative to liso-cel, reflected by covariate effects on Vmax of 0.39 (orva-cel) and 0.20 (ide-cel), corresponding to approximately 1.48-fold and 1.22-fold increases on the original scale (exp(0.39) and exp(0.20), respectively).

Overall, the covariate analysis indicates that, under the same DDE-based structural model, the BCMA-targeted products were characterized by higher baseline levels and higher expansion capacity compared with the CD19-targeted product, while evidence for slower effector-like decay was stronger for orva-cel than for ide-cel.

## Discussion

In prior work, we extended the widely used piecewise CAR-T cellular kinetics framework in a stepwise manner—from the historical constant expansion rate (rho) model to a saturable expansion (Vmax/Km) model, which substantially improved fit and reduced systematic model misspecification during the early expansion phase. ^10, 13, 25^ In the present work, we advanced the Vmax/Km framework further in two complementary directions: (i) replacing discontinuous piecewise switching with smooth S-shaped gating functions to represent a time-varying saturable expansion rate, and (ii) incorporating a delay differential equation (DDE) component to capture a conversion delay (tau1) between effector-like and memory-like compartments (Vmax::r).

Replacing hard switches with smooth gating is primarily a structural/technical refinement, but it has practical value. Abrupt discontinuities can introduce numerical artifacts during estimation and simulation and can complicate interpretation when rate transitions are forced to occur instantaneously. ^14^ In contrast, smooth gating provides a continuous transition that is more stable computationally and better aligned with the biological notion that cellular programs evolve gradually rather than switching ‘on/off’ at a single timepoint. ^26–29^ Importantly, this refinement complements—rather than replaces—the role of DDEs: smooth gating addresses gradual changes in rates, whereas DDEs explicitly represent time lags between upstream processes and downstream outcomes.

The motivation for evaluating DDEs is that time delays are ubiquitous in cellular biology. Signaling cascades, transcriptional reprogramming, commitment to proliferation, cell-cycle progression, and differentiation are not instantaneous. ^21–23^ In T-cell biology, temporal thresholds have been described between stimulation and downstream outcomes such as proliferation commitment and differentiation programs, consistent with the idea that key transitions can be lagged rather than immediate. ^30–33^ DDEs are widely used for representing such lags and have been used broadly in immunodynamics and tumor–immune modeling to represent the time required for proliferation, differentiation, transport, or other intracellular processes. ^21, 34–37^ A related point—also emphasized in the DDE pharmacometrics literature—is that DDEs encode delays at the differential-equation level, which is conceptually distinct from a simple “lag time” added to an observation model. ^38–40^

In our model comparison, the saturable expansion structure (Vmax/Km) provided a substantial improvement over the constant expansion model (rho) across information criteria and diagnostic assessments (Figures 1 and 2), with a clear reduction in bias at the upper end of the observed concentration range (Figure 2a). Incorporating DDE structure yielded additional gains, most notably improving agreement with the line of identity across the mid-range of the data— consistent with a structural enhancement in the model’s ability to reproduce the observed CK profile (Figure 2b). Among the DDE variants evaluated, the most consistent incremental improvement was obtained when the delay was applied specifically to the conversion process (Vmax::r). In contrast, applying delays to effector-like and/or memory-like decay processes (alpha and/or beta), either alone or jointly with conversion (e.g., r/alpha, r/beta, r/alpha/beta), did not yield comparable improvements by ΔAIC/ΔBIC/ΔBICc/ΔOFV (Figures 1 and 2).

Collectively, these results suggest that the data support a lag in the effector to memory conversion process, while additional delayed decay mechanisms were not favored by the model selection criteria. The estimated conversion delay (tau1 ≈ 2.6 days) may reflect the time required for post-activation programming events—such as initiation of a differentiation/commitment program and subsequent phenotypic maturation—before effector-like cells contribute measurably to the memory-like compartment, although tau1 should be interpreted as an empirical kinetic descriptor rather than a direct measurement of any single biological step.

The simulation results provide an intuitive link between structure and inferred compartment dynamics. Moving from rho to Vmax increased predicted expansion-phase magnitude and improved capture of the high end of observed concentrations, consistent with the GOF patterns (Figure 2). The additional step from Vmax to Vmax::r primarily affected the memory-like trajectory, producing higher predicted memory-like cells over a broad time range while leaving the effector-like trajectory comparatively unchanged (Figure 2). The tau1 sensitivity analysis reinforced this interpretation: increasing tau1 shifted the onset/rise/peak timing of the memory-like component later, whereas the effector-like component and total cells showed limited visible sensitivity at the scale shown (Figure 4). This is coherent with the relative compartment magnitudes in our simulations—memory-like cells peak orders of magnitude below the effector-like/total peak—so meaningful changes in the memory-like subpopulation may have minimal impact on total cells in short-term or peak-dominated summaries, yet can still be mechanistically important for longer-term persistence.

Within this framing, tau1 can be positioned as a quantitative descriptor of the timing of memory-like emergence/accumulation under a unified cellular kinetics model, rather than as a purely mathematical artifact. Practically, our results indicate that the dataset is informative for identifying a conversion delay, but provides limited incremental evidence to justify additional delays on decay processes. One plausible explanation is identifiability: expansion and early decline are often richly sampled and strongly inform peak timing and early kinetics, whereas multiple simultaneous delayed decay processes can be difficult to distinguish, especially when they generate partially correlated effects in the terminal phase and when residual variability and sparse late sampling reduce discrimination.

Because Vmax::r provided the best overall fit, we used it as a common structural backbone to compare three CAR-T products spanning two targets (BCMA: orva-cel, ide-cel; CD19: liso-cel) via a categorical product covariate model (Table 2). Under harmonized structural assumptions, the covariate analysis suggested systematic product-level differences in baseline level (conc0), expansion capacity (Vmax), and effector-like decay (alpha). Specifically, both BCMA products exhibited higher baseline concentrations and higher expansion capacity relative to liso-cel, and orva-cel showed evidence for slower effector-like decay. In contrast, the ide-cel alpha effect was small and imprecise (interval included 0), and this uncertainty should be emphasized to avoid overinterpretation. These product-level shifts likely reflect a mixture of factors—target biology, disease context, lymphodepletion and supportive care, manufacturing attributes, and product composition (e.g., CD4/CD8 proportions, memory enrichment)—that cannot be disentangled by a product indicator alone. Accordingly, these findings are best interpreted as empirical differences “as observed” under the analyzed datasets rather than as mechanistic attribution to any single cause.

Two limitations should be acknowledged. First, the effector-like and memory-like compartments are latent constructs; while they yield interpretable trajectories, confirming biological correspondence would require phenotype-resolved measurements (e.g., flow cytometry subsets) and/or integrated links to biomarkers of differentiation state. Second, cross-product comparisons are potentially confounded: a categorical product effect cannot separate product identity from differences in trial design, patient populations, disease burden, inflammation, dose, lymphodepletion, and manufacturing/composition attributes. A natural next step is a structured covariate model incorporating dose and exposure drivers, baseline tumor burden proxies, inflammatory markers, lymphodepletion regimen, and product composition metrics, with explicit attention to collinearity and interpretability.

## Conclusion

A DDE-based CAR-T cellular kinetics model combining saturable expansion, smooth gating, and a conversion delay provided a parsimonious, data-supported description of CAR-T kinetics and outperformed conventional piecewise models based on information criteria and diagnostic evaluations. Simulations indicated that introducing the conversion delay primarily alters the memory-like trajectory, and tau1 sensitivity analyses showed that tau1 mainly controls the timing of the memory-like component with minimal impact on total-cell trajectories at the scale evaluated. Using a unified DDE-based covariate framework, we identified systematic cross-product differences in baseline level and expansion capacity between BCMA- and CD19-targeted therapies, with stronger evidence for slower effector-like decay for orva-cel than for ide-cel. Overall, DDE-based modeling offers a biologically motivated and quantitatively improved framework for CAR-T cellular kinetics and enables consistent cross-product comparisons under harmonized structural assumptions.

## Consent for publication

All the authors have reviewed and concurred with the manuscript.

## Authors’ contributions

Y.C. and Y.L. contributed to conception and design; Y.C. and Y.L. contributed to acquisition of data; Y.C. and Y.L. contributed to analysis; all authors contributed to interpretation of data; Y.C. and Y.L. drafted and revised the article. Both authors made substantial contributions to conception and design, acquisition of data, or analysis and interpretation of data; took part in drafting the article or revising it critically for important intellectual content; agreed to submit to the current journal; gave final approval of the version to be published; and agreed to be accountable for all aspects of the work.

---

**Supplementary Figure 1.**
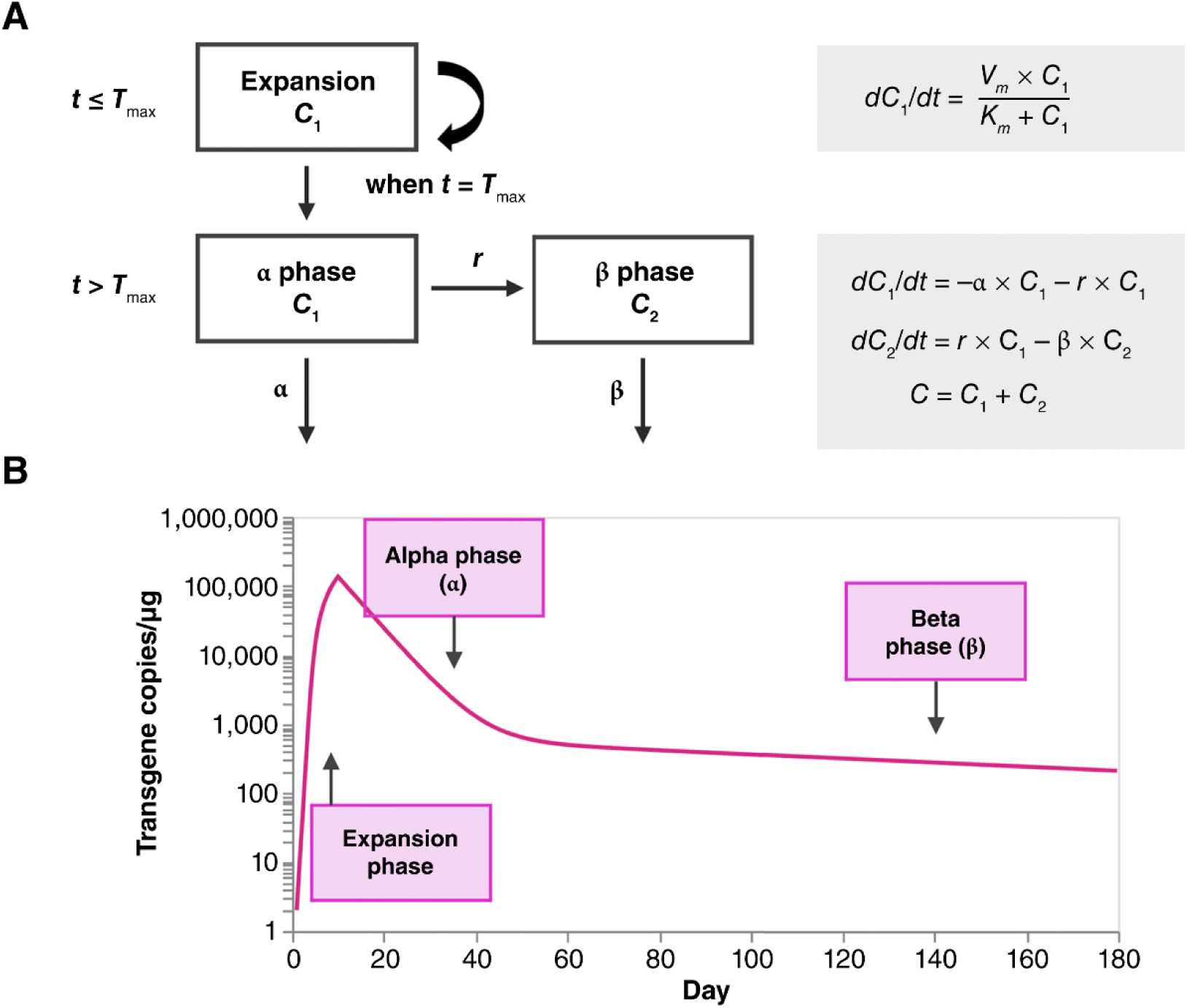
Semi-mechanistic piecewise saturable expansion model for CAR-T cellular kinetics. (**A)** Compartmental model and the associated differential equations that describe T-cell kinetics during the expansion and contraction phases. (**B)** Graphical representation of the typical cellular kinetics profile. *C*_1_, transgene levels in expansion phase (time < *T*_max_) or α decay phase (time ≥ *T*_max_); *C*_2_, transgene levels in β decay phase; *Km*, transgene levels giving expansion rate of 1/2 *Vm*; *r*, conversion rate constant from rapid contraction phase to slow contraction phase; *T*_max_, time to maximum transgene level; *Vm*, maximum expansion rate; α, rate constant of rapid contraction phase; β, rate constant of slow contraction phase.

